# Coordinated IL-2 and TGF-β Signaling Activates Stable Regulatory T Cells and Promotes Immune Tolerance

**DOI:** 10.64898/2026.03.09.709648

**Authors:** Nicholas M. Jackson, Javier A. Carrero, Kristin Yeung, Evelyn Needham, Emily Christie, Natalie Yakobian, Challen Pretorius, Stella Hoft, Kirsten Foley, Eric Ford, Simon Low, Ronald Seidel, Anish Suri, Natasha M. Girgis, Steven N. Quayle, Richard J. DiPaolo

## Abstract

Establishing immune tolerance requires the integration of cytokine signals that favor regulatory T cell (Treg) function over effector differentiation, yet how such coordination can be achieved selectively in vivo remains incompletely defined. Regulatory T cells are essential for maintaining immune homeostasis and peripheral tolerance, and both interleukin-2 (IL-2) and transforming growth factor–β (TGF-β) contribute to Treg activation, differentiation, and stability in a highly context-dependent manner. Here, we examine the consequences of coordinated IL-2 and TGF-β3 signaling using an engineered cytokine construct combining attenuated IL-2 with receptor-masked TGF-β3. In vitro, attenuated IL-2/TGF-β3 signaling promoted conversion of naïve CD4⁺ T cells into Foxp3⁺ induced Tregs, expanded endogenous Tregs, and suppressed inflammatory cytokine production by memory and effector CD4⁺ T cells. In vivo, a single administration selectively increased the frequency of CD4⁺Foxp3⁺ regulatory T cells exhibiting an activated and stable phenotype, while minimally activating Foxp3⁻ conventional CD4⁺ T cells relative to IL-2 alone. In a Treg-deficiency–driven adoptive transfer model of autoimmunity, early exposure to coordinated IL-2/TGF-β3 signaling enhanced Treg activation, reduced tissue infiltration by autoreactive T cells, and conferred sustained protection from autoimmune gastritis in the absence of continued treatment. Together, these findings identify a mode of coordinated IL-2 and TGF-β3 signaling sufficient to stabilize regulatory T cell responses and promote immune tolerance while limiting activation of conventional CD4⁺ T cells, highlighting signal integration as a determinant of tolerogenic immune regulation in vivo.

**One Sentence Summary:** CUE-401 coordinates IL-2 and TGF-β signaling to expand and activate regulatory T cells and restrain autoimmune responses.

## Introduction

Autoimmune diseases arise from the breakdown of peripheral immune tolerance, resulting in persistent immune-mediated damage to self-tissues. The discovery of CD4⁺CD25⁺ regulatory T cells (Tregs) marked a paradigm shift in understanding immune tolerance (*1*). Tregs maintain tolerance by suppressing autoreactive effector T cell responses while preserving protective immunity through multiple mechanisms, including consumption of interleukin-2 (IL-2), expression of inhibitory receptors (e.g. CTLA-4), and production of immunoregulatory cytokines, such as transforming growth factor–β (TGF-β) (*2*). The essential role of Tregs in immune homeostasis is underscored by the severe autoimmune pathology observed in humans and mice with mutations in the transcription factor Foxp3, which is required for Treg lineage commitment and stability (*3–6*).

Tregs are classically defined by expression of CD4, the high-affinity IL-2 receptor α-chain (CD25), and Foxp3. This population includes thymus-derived natural Tregs (nTregs) and peripherally induced Tregs (pTregs), which arise from conventional CD4⁺ T cells in response to antigenic stimulation and cytokine cues (*7, 8*). In vitro, naïve CD4^+^ T cells stimulated with TCR signals plus TGF-β, and IL-2 differentiate into Foxp3⁺ induced Tregs (iTregs) with potent suppressive activity (*9*). Precise modulation of IL-2– dependent STAT5 activation together with TGF-β–mediated Foxp3 induction is also critical for maintaining stable, suppressive Tregs (*9–11*). Accordingly, this cytokine combination has been widely used to generate suppressive iTregs ex vivo for adoptive cell therapies that are effective in preclinical models of autoimmunity and graft-versus-host disease (*12–14*).

The immunosuppressive capacity of Tregs has motivated their therapeutic use in transplantation and immune-mediated diseases. Several different IL-2 variants (muteins) have been engineered to activate and expand Tregs and are being tested for effectiveness in treating inflammatory diseases (*15–17*). Key challenges include a limited therapeutic window due to off-target stimulation of non-Treg populations, the short half-life of IL-2, and the need for precise dosing to achieve selective Treg engagement. Moreover, Treg induction, stability, and specialization in vivo often require additional cues, including TGF-β.

Coordinating IL-2 and TGF-β signaling in vivo is challenging due to short in vivo half-lives of cytokines, tolerability considerations, and the need to co-engage receptors on the same target cell(s). CUE-401, a novel Fc-based IL-2/TGF-β3 fusion protein designed for coordinated delivery of IL-2 and TGF-β3 signals in vivo, was designed to address these challenges. CUE-401 includes an attenuated IL-2 mutein (F42A, H16A) with reduced affinity for CD25 and CD122 (*18–20*), and a novel receptor-masked attenuated TGF-β3 variant, linked via flexible linkers to an Fc scaffold to promote IL-2/TGF-β3 co-signaling and extend half-life. All amino acid sequences are human-derived, and both cytokines cross-react with mouse receptors (*21, 22*). Here, we evaluated the biological activities of CUE-401 in vitro and in vivo for its ability to enhance Tregs, suppress effector T cell responses, and mitigate autoimmune pathology.

## Results

### CUE-401 delivers IL-2 and TGF-β signals to T cells

CUE-401 is a bispecific fusion protein composed of affinity attenuated variants of IL-2 and TGF-β3 (Fig. 1A). The IL-2 mutein (F42A and H16A) reduces affinity to IL-2Rα (CD25) and IL-2Rβ (CD122) (*18*). The TGF-β3 component is engineered as a monomeric, receptor masked variant to attenuate receptor affinity. The TGF-β3 moiety carries a C77S mutation that prevents formation of the interchain disulfide bond while maintaining its ability to assemble the natural signaling complex. Masking of the TGF-β3 moiety is achieved through fusion to a modified TGF-β receptor II extracellular domain lacking the N-terminal residues required for productive association with TGF-β receptor I and containing a D118A substitution that reduces affinity for TGF-β3 relative to the wild-type receptor (*23, 24*). This configuration limits constitutive TGF-β receptor engagement while permitting biological activity upon cell-associated presentation. The IL-2 and TGF-β3 domains are connected via flexible linkers to a heterodimeric Fc scaffold to support coordinated signaling and molecular stability.

**Fig. 1.**
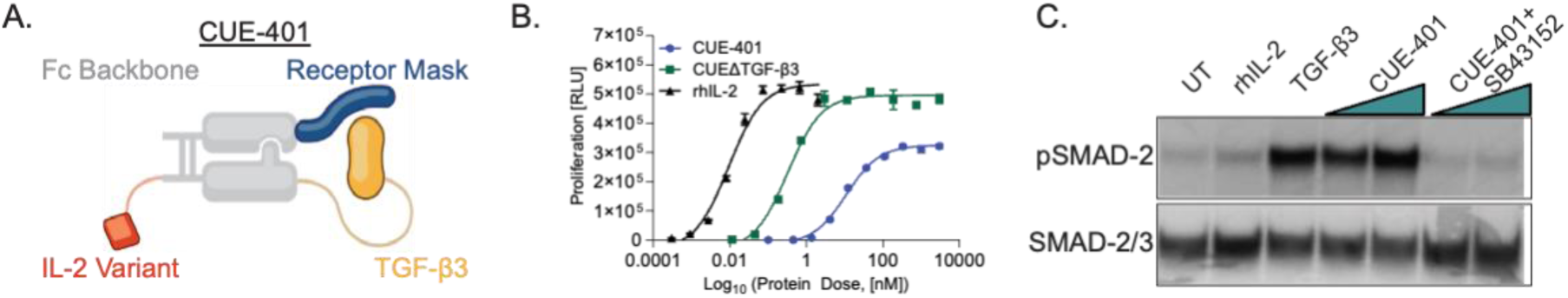
CUE-401 promotes IL-2 and TGF-β signaling activity. (**A**) Model of CUE-401 design. (**B**) CTLL-2 proliferation with recombinant human IL-2 (Proleukin), CUE-401, or CUE-401ΔTGF-β3. (**C**) Immunoblot detecting phosphorylated SMAD-2 (pSMAD-2) and total SMAD-2/3 after 30 minutes of stimulation with rhIL-2, rhTGF-β3, CUE-401, or CUE-401 + SB43152.

To quantify IL-2 activity, CTLL-2 proliferation assays were performed comparing wild-type human IL-2 (Proleukin), CUE-401, and a construct containing the attenuated IL-2 moiety lacking the TGF-β3 domain (CUE-401ΔTGFβ; Fig. 1B). Both CUE-401 and CUE-401ΔTGFβ induced dose-dependent proliferation that was reduced compared to Proleukin, consistent with attenuated IL-2 receptor signaling (Fig. 1B). TGF-β3 signaling activity was assessed by SMAD2/3 phosphorylation in murine splenocytes. CUE-401 induced robust phosphorylation of SMAD2/3 comparable to recombinant TGF-β3, whereas recombinant IL-2 alone failed to do so. SMAD2/3 phosphorylation induced by CUE-401 was abrogated by pretreatment with the selective TGF-β receptor inhibitor SB431542, confirming engagement of canonical TGF-β receptor signaling pathways (Fig. 1C).

### CUE-401 induces and expands Tregs and suppresses effector T cell function in vitro

Having established that both cytokine components of the engineered construct were biologically active, we next examined whether coordinated IL-2 and TGF-β3 attenuated signaling could induce de novo Foxp3 expression in CD4⁺ T cells. CD4⁺CD8^-^Foxp3.GFP^-^ thymic T cells were sorted from thymi of BALB/c.Foxp3.eGFP mice and stimulated with anti-CD3/anti-CD28 in the presence of recombinant human IL-2 (rhIL-2), rhIL-2 plus recombinant human TGF-β3 (rhTGF-β3), or the engineered IL-2/TGF-β3 construct (Fig. 2). Stimulation with IL-2 alone induced minimal Foxp3 expression (∼1%), whereas combined IL-2 and TGF-β3 signaling, delivered either as separate recombinant cytokines or engineered attenuated IL-2/TGF-β-Fc configuration, induced Foxp3 expression in more than 75% of CD4⁺ T cells (Fig. 2A). Foxp3 induction occurred in a dose-dependent manner, with an EC₅₀ of ∼18 nM (Fig. 2B). These findings indicate that coordinated and attenuated IL-2/TGF-β3 signaling is sufficient to drive inducible regulatory T cell (iTreg) differentiation from naïve CD4⁺ T cells.

**Fig. 2.**
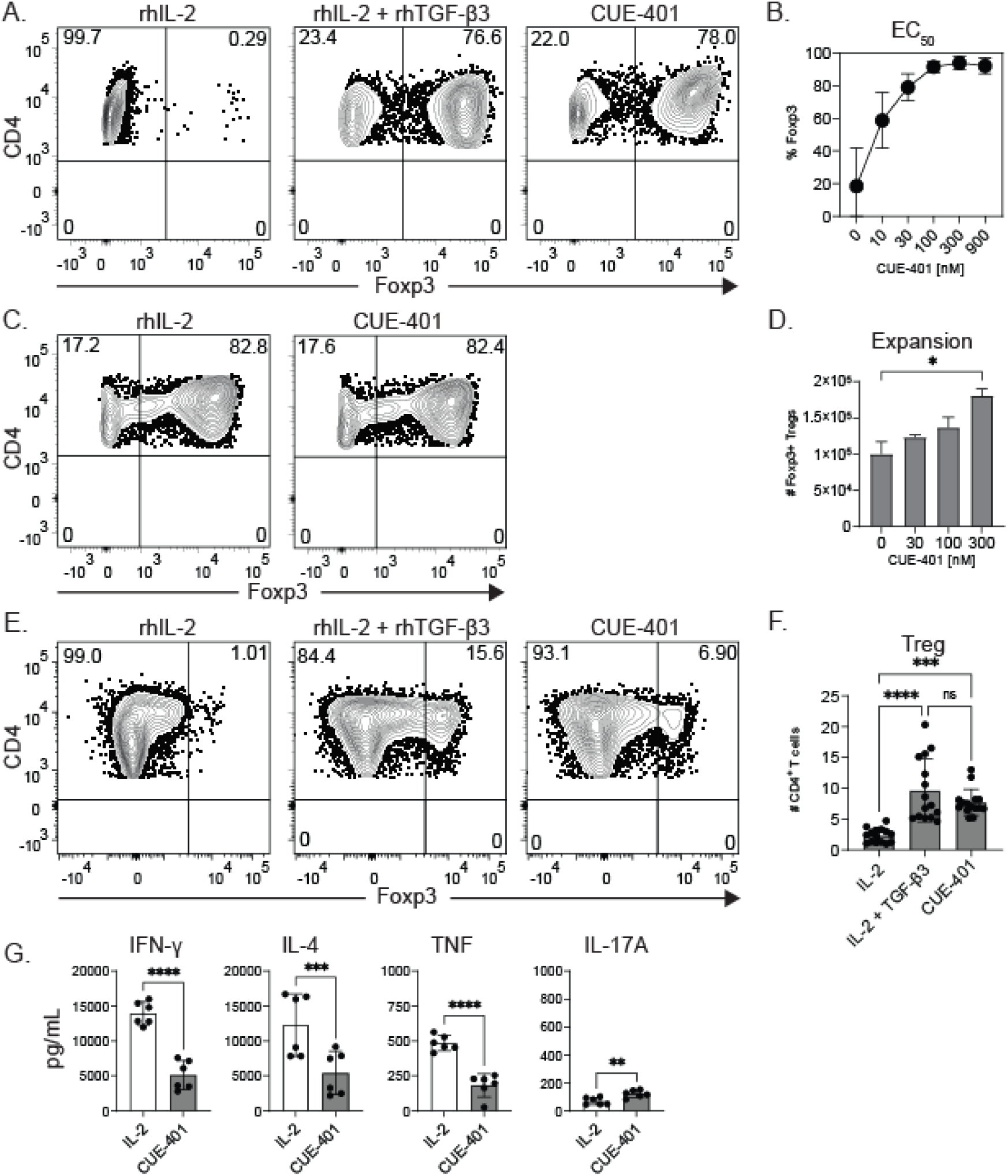
CUE-401 promotes iTreg induction, nTreg expansion, and Teff suppression. (**A**). Representative flow cytometric analysis showing the percentage of CD4^+^ gated cells expressing Foxp3 in vitro following stimulation with rhIL-2, rhIL-2 + rhTGF-β3, or CUE-401. (**B**). Dose-response curve displaying Foxp3 expression with increasing doses of CUE-401. (**C**). Representative flow cytometric analysis showing the percentage of CD4^+^ gated cells expressing Foxp3 after six days in vitro stimulation of Tregs with rhIL-2 or CUE-401. (**D**). Number of Foxp3 expressing cells at various concentrations of CUE-401. (**E**). Flow cytometric analysis displaying the percentage of CD4^+^ gated cells expressing Foxp3 after six days in vitro simulation of Teff with rhIL-2, rhIL-2 + rhTGF-β3, or CUE-401. (**F**) Percentages of CD4^+^Foxp3^+^ Tregs in rhIL-2, rhIL-2 + rhTGF-β3 and CUE-401 treated samples. (**G**). Amounts of IFN-γ, IL-4, TNF, and IL-17A secreted from CD4^+^CD44^hi^ Foxp3.GFP^-^ T cells with rhIL-2 or CUE-401 treatment. Flow cytometry plots are representative of individual sorted samples analyzed from thymus (A) or spleen (D,F,G) of 4-8 healthy BALB/c mice. EC_50_ calculated by taking the concentration halfway between the top and bottom value (approximately 20-80%). Bars in each graph represent the mean +/-SD. *P*-values calculated using Kruskal-Wallis test (D), ordinary one-way ANOVA with Tukey’s multiple comparisons test (F), or Wilcoxon matched-pairs signed rank test (G) with \**p* = 0.0332, \*\**p* = 0.0021, \*\*\**p* = 0.0002, and \*\*\*\**p* < 0.0001.

To assess whether CUE-401 expands endogenous CD4^+^Foxp3^+^ Tregs, CD4⁺Foxp3.eGFP⁺ Tregs sorted from the spleens of BALB/c.Foxp3.eGFP mice were stimulated with anti-CD3/anti-CD28 in the presence of presence of either rhIL-2 alone or the engineered IL-2/TGF-β3 construct. As expected, Tregs maintained Foxp3 expression when cultured with rhIL-2 (Fig. 2C). Similarly, Tregs stimulated in the presence of CUE-401 maintained Foxp3 expression and expanded in a dose-dependent manner (Fig. 2D), indicating that this signaling configuration supports both lineage stability and proliferative capacity of mature Tregs.

To determine the effects of coordinated signaling of the attenuated IL-2/TGF-β3 influences effector/memory T cells, CD4⁺CD44^hi^Foxp3.eGFP⁻ cells sorted from spleens of BALB/c.Foxp3.eGFP mice were stimulated with anti-CD3/anti-CD28 in the presence of either rhIL-2, rhIL-2 + rhTGF-β3, or CUE-401 (Fig. 2E). Treatment with rhIL-2 resulted in minimal induction of Foxp3 expression (∼1%). However, stimulation of these purified effector/memory T cells in the presence of rhIL-2 + rhTGF-β3 or CUE-401 induced Foxp3 expression in ∼10% of effector/memory CD4⁺ T cells (Fig. 2E-F). We further assessed cytokine production by effector/memory CD4⁺ T cells under these conditions. Stimulation in the presence of IL-2 alone resulted in robust secretion of IFN-γ, IL-4, TNF, and IL-17A. In contrast, coordinated IL-2/TGF-β signaling markedly reduced secretion of IFN-γ, IL-4, and TNF relative to IL-2 alone (Fig. 2G), indicating suppression of pro-inflammatory effector function.

Collectively, these results define three distinct immunological outcomes of the coordinated signaling by attenuated the IL-2 and TGF-β signaling in vitro: induction of Foxp3⁺ regulatory T cells from naïve CD4⁺ T cell precursors, maintenance and expansion of endogenous regulatory T cells, and partial reprogramming of effector/memory CD4⁺ T cells toward a less inflammatory state.

### Administration of coordinated IL-2/TGF-β induces robust expansion of stable regulatory T cells in mice

To assess the in vivo consequences of coordinated signaling by attenuated IL-2 and TGF-β, BALB/c mice received a single administration of the engineered cytokine construct or vehicle control (PBS). Six days after treatment, the frequency of CD4⁺Foxp3⁺ regulatory T cells in the spleen increased to approximately 61% of the CD4⁺ T cell compartment in treated animals, compared with approximately 20% in control mice (Fig. 3A-B). To determine whether regulatory T cells expanded under these conditions exhibited features of lineage stability, we analyzed methylation of the Treg-specific demethylated region (TSDR), a CpG-rich enhancer element within the Foxp3 locus (CNS2) associated with stable Foxp3 expression (*25*). Splenocytes from vehicle-treated and cytokine-treated mice were sorted into CD4⁺Foxp3.eGFP⁺ and CD4⁺Foxp3.eGFP⁻ populations (Fig. 3C). Regulatory T cells from treated mice displayed a degree of TSDR demethylation comparable to that observed in endogenous regulatory T cells from control animals, whereas conventional CD4⁺Foxp3⁻ T cells from treated mice retained a methylation profile indistinguishable from that of control conventional T cells (Fig. 3D). Consistent with these epigenetic features, regulatory T cells isolated from both treated and control mice maintained Foxp3 expression following ex vivo restimulation for six days in the absence of continued cytokine exposure (Fig. 3E). Together, these findings indicate that coordinated the IL-2 and TGF-β signaling provided by CUE-401 selectively expands regulatory T cells in vivo while preserving epigenetic and transcriptional features associated with stable lineage commitment.

**Fig. 3.**
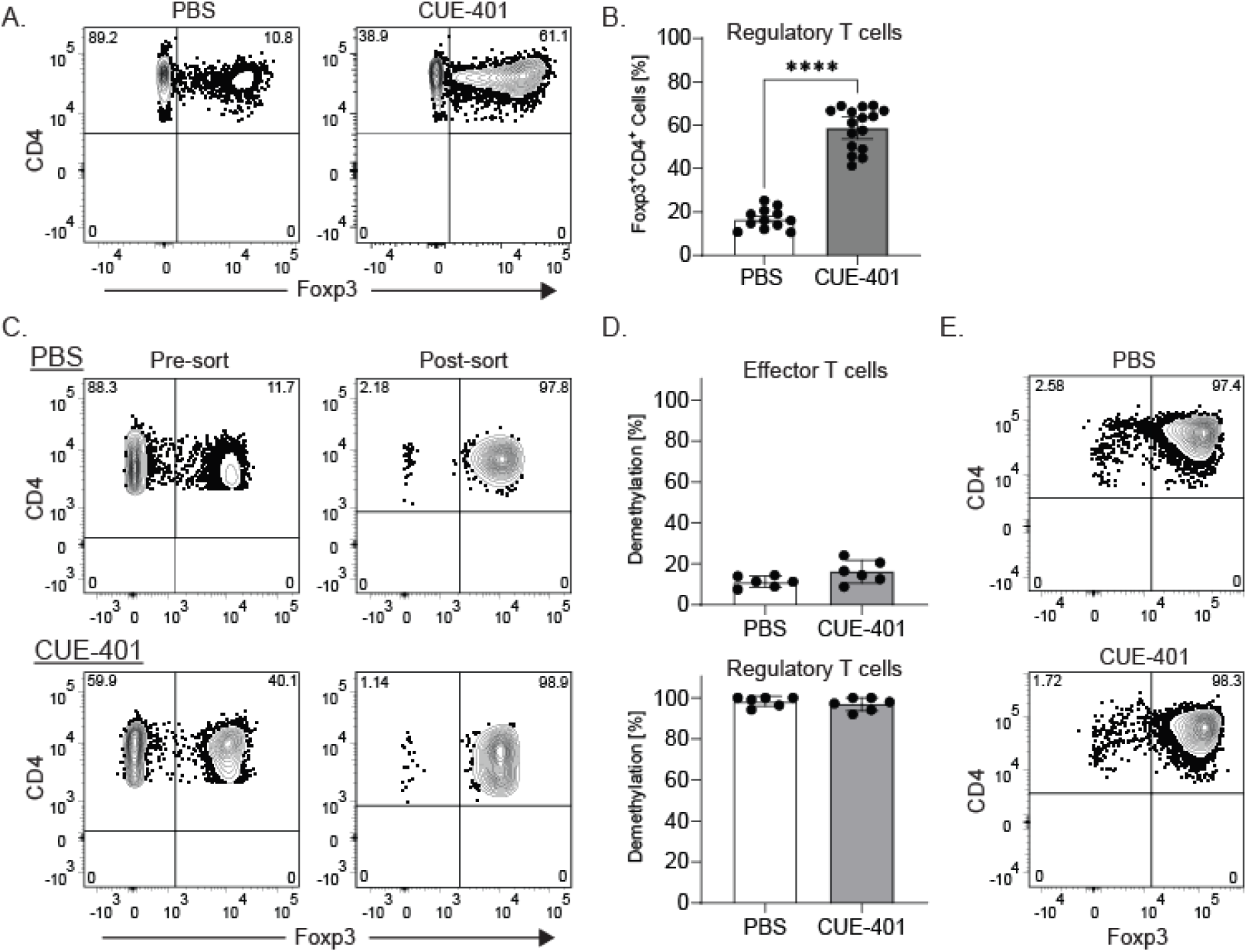
CUE-401 treatment increases proportions of stable Foxp3+ Tregs in vivo. (**A**) Flow cytometric analysis showing percentages of CD4^+^ gated cells expressing Foxp3 six days after treatment of BALB/c mice. (**B**) Combined data for the percentages of CD4^+^Foxp3^+^ Tregs in PBS and CUE-401 treated mice. (**C**) Flow cytometric analysis of CD4^+^Foxp3^+^ expression prior to and after performing FACS to purify CD4^+^Foxp3^+^GFP^+^ Tregs. (**D**) Results of qPCR quantifying the relative percentage of DNA demethylation at key CpG sites within the Treg-specific demethylated region (TSDR) of the *Foxp3* locus of CD4^+^Foxp3.eGFP^-^ and CD4^+^Foxp3.eGFP^+^ cells sorted from PBS and CUE-401 treated mice. (**E**) Representative flow cytometric plots showing Foxp3 expression of purified T cells from PBS and CUE-401 treated mice after 6 days of restimulation in vitro in the absence of cytokines. For all graphs, controls were PBS treated age-matched mice. Data in (A-B) was generated from spleens of 2-5 mice in 7 independent experiments. Plots in (C) show the average proportion of Foxp3^+^ T cells pre and post sorting from 8 mice in 2 independent experiments. Bars represent the median +/- 95% C.I. TSDR analysis in (D) includes 12 mice from 2 independent experiments. Female mice methylation status was normalized for methylation state on both x chromosomes. *P* values were calculated utilizing the Mann-Whitney *U* test. \*\*\*\**p* < 0.001.

### Expanded regulatory T cells exhibit an activated phenotype following combined IL-2/TGF-β administration

Having established that coordinated IL-2 and TGF-β3 signaling induces robust expansion of regulatory T cells in vivo, we next examined the phenotypic characteristics of regulatory T cells six days following treatment. Consistent with earlier observations, treatment resulted in a marked increase in the frequency of CD4⁺Foxp3⁺ regulatory T cells relative to vehicle-treated controls (Fig. 4A). We therefore assessed expression of surface and intracellular markers associated with regulatory T cell activation and suppressive function, including Foxp3, GITR, CD25, CTLA-4, CD103, CD73, CD38, CD134, CD137, and Ki67 (*26, 27*). Multidimensional flow cytometry analysis followed by UMAP projection revealed enrichment of regulatory T cell clusters exhibiting features of activation in treated mice, including increased frequencies of GITR^hi^/CD25^hi^ and CTLA-4^hi^/CD103^hi^ Tregs, and higher expression of CD25, CTLA-4, and GITR overall (Fig. 4A–C). At the population level, regulatory T cells from treated animals displayed increased expression of CD25, CTLA-4, and GITR, consistent with enhanced responsiveness to IL-2 and acquisition of suppressive effector features (Fig. 4C).

**Fig. 4.**
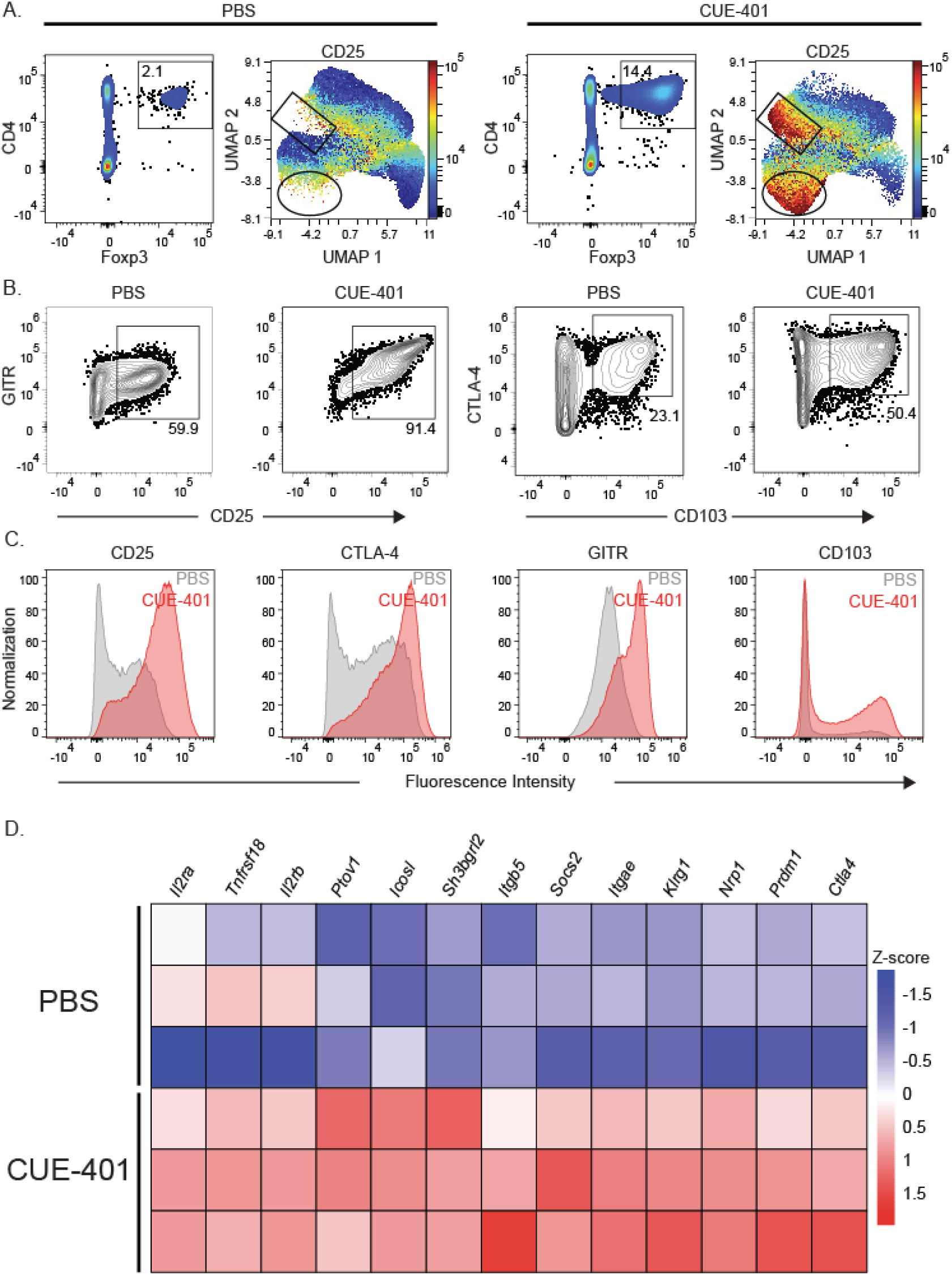
CUE-401 increases expression of transcripts and proteins associated with Treg activation. (**A**). Flow cytometric analysis shows the percentage of CD4^+^Foxp3^+^ expressing cells following treatment of mice with PBS or CUE-401. CD4⁺Foxp3⁺ Tregs were identified, and UMAP projections were generated using OMIQ software to cluster cells based on protein expression. The feature plot shows CD25 expression within the Foxp3⁺ populations. (**B**). Cells were gated on the CD4⁺Foxp3⁺ populations to assess surface expression of GITR, CD25, CTLA-4 and CD103. (**C**). Histograms display the expression of GITR, CD25, CTLA-4, and CD103 within the Treg compartment of PBS-treated controls compared to CUE-401–treated mice. (**D**). Heatmap from bulk RNA sequencing of sorted Tregs, highlighting differentially expressed genes between PBS and CUE-401 treated mice. Genes included had a log₂ fold change > 2 and adjusted p-value < 0.05, using the Benjamini–Hochberg correction to control false discovery rate (FDR). For all flow cytometry plots, data are representative of individual spleens from a single experiment with 3 mice per treatment group. PBS-treated, age-matched mice served as controls in all analyses.

To further characterize the transcriptional state of regulatory T cells expanded under coordinated IL-2/TGF-β3 signaling, we performed bulk RNA sequencing on purified CD4⁺Foxp3⁺ T cells from treated and control mice (Fig. 4D). Regulatory T cells from treated animals exhibited increased expression of genes associated with regulatory T cell activation, tissue localization, and suppressive function, including *Itgae* (CD103), *Nrp1*, *Il2ra*, *Il2rb*, *Lag3*, *Tnfrsf18* (GITR), *Icosl*, and *Klrg1* (*28–32*). Together, these phenotypic and transcriptional analyses indicate that regulatory T cells expanded under coordinated IL-2 and TGF-β signaling acquire an activated regulatory program associated with enhanced suppressive capacity, complementing the observed epigenetic and transcriptional stability of the expanded population (*33–36*).

### Combined IL-2 and TGF-β signaling responses in Tregs are distinct from IL-2 alone

To isolate the contribution of TGF-β signaling and distinguish the biological effects of coordinated IL-2/TGF-β3 signaling from those of IL-2 alone, we compared the engineered IL-2/TGF-β3 construct (CUE-401) with an otherwise identical IL-2–only variant lacking the TGF-β3 domain (CUE-401ΔTGF- β3). Both constructs incorporate the same attenuated IL-2-Fc moiety, enabling direct comparison of IL-2 signaling in the presence or absence of concurrent TGF-β3 input. In vivo, administration of either construct resulted in significant expansion of splenic regulatory T cells; however, regulatory T cell frequencies were consistently higher following coordinated IL-2/TGF-β3 signaling (Fig. 5A, B). Moreover, regulatory T cells expanded under coordinated IL-2/TGF-β3 signaling conditions exhibited a greater proportion of GITR^hi^ CD38^hi^ Tregs compared with those expanded by IL-2 alone (Fig. 5C, D), consistent with acquisition of an activated regulatory phenotype.

**Fig. 5.**
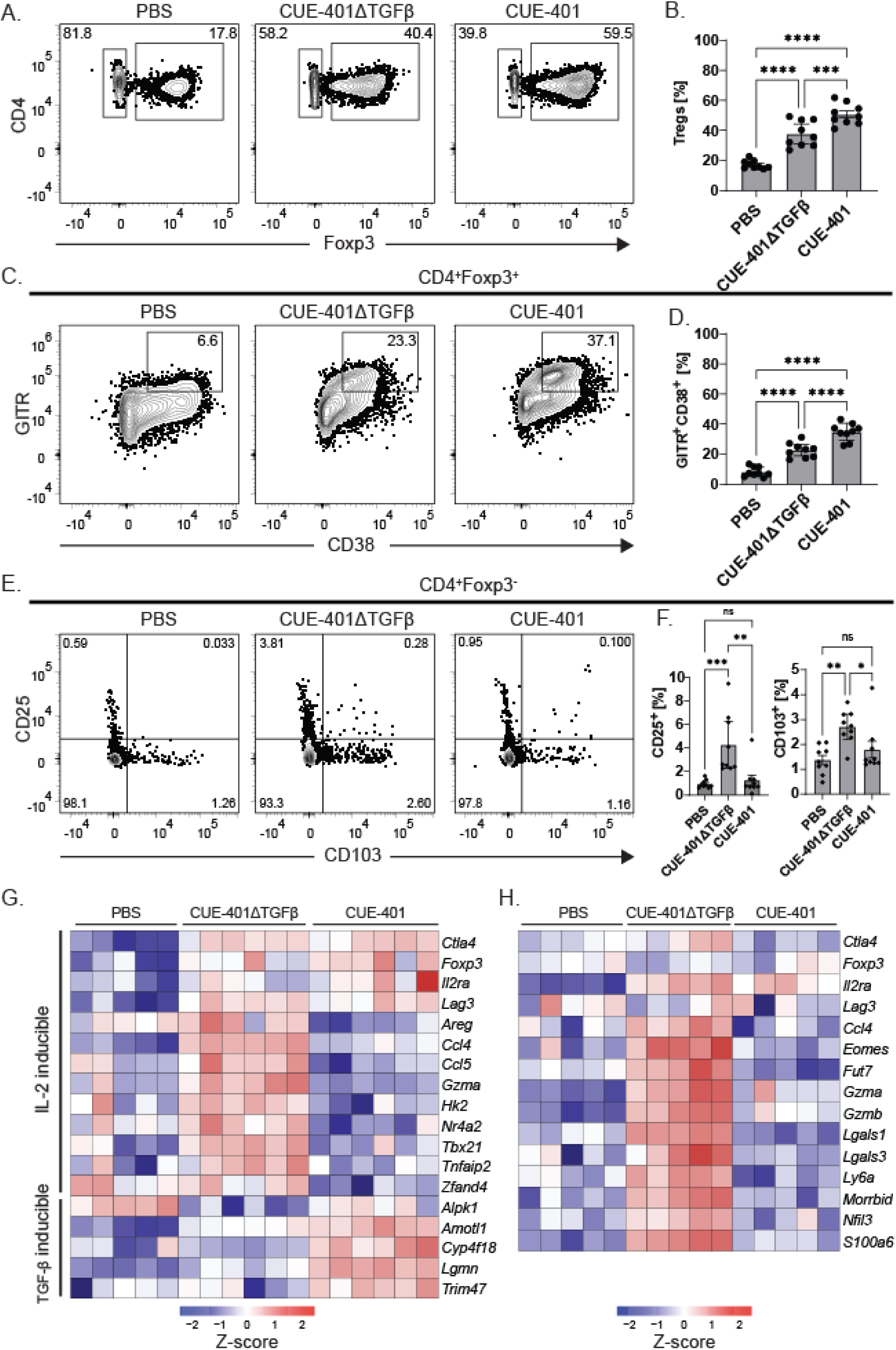
The combined effects of IL-2 and TGF-β3 in CUE-401 differ from IL-2 treatment alone. **(A)**. Representative flow cytometric analysis showing the percentage of CD4^+^ gated cells expressing Foxp3 six days after treatment with PBS (left), CUE-401ΔTGF-β3 (middle), or CUE-401 (right) in vivo. **(B)**. Combined data showing the percentages of CD4^+^Foxp3^+^ Tregs in PBS, CUE-401ΔTGF-β3, or CUE-401 treated mice. (**C**). Representative flow cytometric analysis comparing GITR by CD38 expression in CD4^+^Foxp3^+^ Tregs following the indicated treatments. (**D**). Combined data showing the percentage of GITR^+^CD38^+^ Tregs following PBS, CUE-401ΔTGF-β3, or CUE-401 treatment. (**E**). Representative flow cytometric analysis comparing CD25 by CD103 of CD4^+^Foxp3^-^ gated cells. (**F**). Combined data showing the percentage of CD25^+^CD4^+^Foxp3^-^ cells and CD103^+^CD4^+^Foxp3^-^ cells six days after treatment. Data shown are representative of two independent experiments (N = 9 per group). Bars represent the median ± 95% confidence interval. (**G**, **H**). Heatmaps of gene expression profiles in sorted CD4⁺Foxp3.eGFP⁺ Treg cells or CD4^+^Foxp3.eGFP^-^ Tconv cells, derived from a subset of control or treated mice (N = 2,3 per group, respectively). *P* values for flow results were determined utilizing an ordinary one-way ANOVA with Tukey’s multiple comparisons test with \**p* = 0.0332, \*\**p* = 0.0021, *** *p* = 0.0002, and \*\*\*\**p* < 0.0001.

Notably, IL-2 signaling in the absence of TGF-β3 led to broader activation of conventional CD4⁺ T cells. (Fig. 5E–F). Treatment with the IL-2–only construct increased expression of CD25 and CD103 on Foxp3⁻ conventional CD4⁺ T cells, whereas coordinated IL-2/TGF-β3 signaling did not induce comparable upregulation of these markers in the conventional T cell compartment (Fig. 5E–F), indicating enhanced regulatory selectivity when TGF-β signaling is integrated with IL-2.

To further define differences in signaling outcomes at the transcriptional level, we performed RNA sequencing on sorted CD4⁺Foxp3.eGFP⁺ regulatory T cells and CD4⁺Foxp3.eGFP⁻ conventional T cells from mice treated with vehicle, IL-2 alone (CUE-401ΔTGF-β3), or coordinated IL-2/TGF-β3 signaling (CUE-401). Both IL-2 alone and coordinated IL-2/TGF-β signaling increased expression of core regulatory T cell–associated genes, including *Ctla4*, *Foxp3*, *Il2ra*, and *Lag3*, relative to vehicle-treated control (Fig. 5G). However, several transcripts induced in regulatory T cells by IL-2 alone were not comparably induced when IL-2 signaling occurred in conjunction with TGF-β3, indicating qualitative differences in downstream signaling programs. Conversely, coordinated IL-2/TGF-β3 signaling selectively induced a set of transcripts previously associated with TGF-β–dependent regulation, including *Alpk1*, *Amot1*, *Cyp4f18*, *Lgmn*, and *Trim47*, which were not induced by IL-2 alone. Together, these findings demonstrate that integrated IL-2 and TGF-β signaling establishes a regulatory T cell transcriptional state that is distinct from, and not merely an amplification of, the program elicited by IL-2 in isolation.

In contrast, within the CD4⁺Foxp3⁻ conventional T cell compartment, IL-2 signaling in the absence of concurrent TGF-β input elicited a transcriptional program characteristic of IL-2–driven cellular activation. Treatment with the IL-2–only construct (CUE-401ΔTGF-β3) resulted in increased expression of genes associated with effector differentiation and activation, including *Ccl4*, *Eomes*, *Fut7*, *Gzma*, *Gzmb*, *Lgals1*, *Lgals3*, *Ly6a*, *Morrbid*, *Nfil3*, and *S100a6*. These transcripts were not comparably induced in conventional CD4⁺ T cells from mice receiving coordinated IL-2 and TGF-β3 signaling (CUE-401, Fig. 5H). Together, these data indicate that IL-2 signaling alone promotes activation-associated transcriptional programs in conventional CD4⁺ T cells, whereas integration of TGF-β3 signaling constrains this response. Thus, coordinated IL-2 and TGF-β3 signaling differentially regulates regulatory and conventional CD4⁺ T cell compartments, favoring regulatory T cell–specific activation while limiting IL-2–driven activation of conventional T cells.

### Early coordinated IL-2 and TGF-β signaling suppresses autoreactive T cell responses and limits autoimmune gastritis

To examine the consequences of coordinated IL-2 and TGF-β3 signaling in an established model of Treg deficiency–driven autoimmunity, we employed a well-characterized murine model of autoimmune gastritis in which CD25⁺ regulatory T cells are depleted from BALB/c splenocytes prior to transfer into athymic BALB/c *nu/nu* recipients (*1*). Depletion of regulatory T cells in this system reliably results in the development of autoimmune gastritis in all recipients. To enable tracking of antigen-specific autoreactive conventional CD4⁺ T cells, a small number of CD4⁺ TCR-transgenic T cells specific for the gastric autoantigen H⁺/K⁺ ATPase and expressing the Thy1.1 congenic marker (*37*) were admixed with CD25-depleted BALB/c splenocytes (Thy1.2⁺) before transfer. A representative flow cytometric analysis of the transferred CD4⁺ T cell populations is shown in Fig. 6A.

**Fig. 6.**
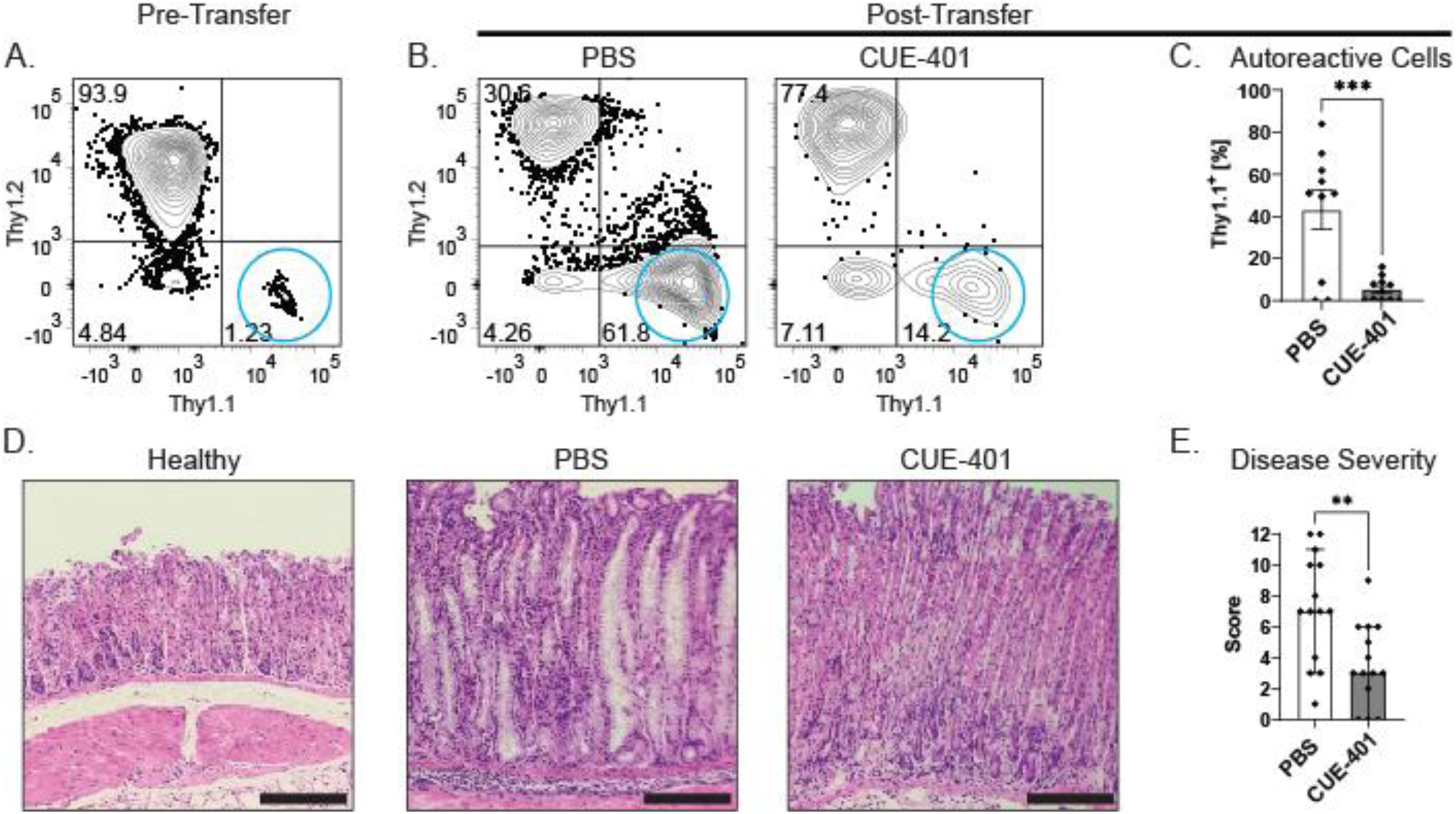
CUE-401 treatment inhibits Teff responses and decreases autoimmune disease severity. **(A)**. Flow cytometric analysis of Thy1.2^+^CD4^+^ splenocytes and autoreactive Thy1.1^+^CD4^+^ splenocytes prior to injection into BALB/c *nu/nu* mice. (**B**). Flow cytometric analysis of Thy1.1 and Thy1.2 expression in CD4^+^ T cells from stomachs of BALB/c *nu/nu* mice ∼62 days after transfer. (**C**) Combined analysis of Thy1.1 expressing autoreactive cells in the gastric mucosa of nude mice ∼62 days after transfer and subsequent treatment with PBS or CUE-401. Quantified data are presented as mean ± SEM. Experiments were conducted independently three times, with 2–5 mice per group per experiment. (**D**). Representative images from four independent experiments, comparing tissues from healthy untreated BALB/c mice, PBS-treated BALB/c *nu/nu* controls, and CUE-401–treated BALB/c *nu/nu* mice. (**E**). Histological disease scores were determined based on multiple pathological features, including inflammation, glandular atrophy, and epithelial hyperplasia. The final score represents the combined values from these categories. Graph bars represent the median +/- 95% C.I. Experiments contained 3-5 aged matched female mice. *P*-values were determined utilizing either Unpaired Student’s T test (C) or Mann-Whitney *U* test (D) with \*\**p* = 0.0040 and \*\*\**p* = 0.0005. Scale bars in micrographs represent 180 µM.

Recipient BALB/c *nu/nu* mice were administered vehicle or CUE-401 on days 1 and 14 following transfer. Sixty-two days after transfer, mice receiving CUE-401 exhibited a marked reduction in the frequency of autoreactive H⁺/K⁺ ATPase-specific CD4⁺Thy1.1⁺ T cells infiltrating the gastric mucosa compared with vehicle-treated controls (approximately 10% versus 44%, respectively; *P* = 0.0005; Fig. 6B–C). Histopathological examination of stomach tissue was performed to assess disease severity, including inflammation, glandular atrophy, and epithelial hyperplasia, hallmark features of autoimmune gastritis (*38*). As expected, vehicle-treated mice developed moderate-to-severe autoimmune gastritis, with a median disease score of 6 on a 12-point scale. In contrast, mice that received coordinated IL-2/TGF-β3 signaling displayed significantly reduced pathology, with a median disease score of 3, and no detectable gastritis in a subset of animals (Fig. 6D, E). Together, these findings indicate that early and transient exposure to coordinated IL-2 and TGF-β3 signaling is sufficient to durably constrain autoreactive CD4⁺ T cell accumulation and limit tissue pathology in a stringent model of regulatory T cell deficiency–driven autoimmunity.

### Coordinated IL-2 and TGF-β signaling induces a distinct regulatory T cell state in a disease context

To investigate the immunological mechanisms underlying suppression of autoimmune gastritis following coordinated IL-2 and TGF-β3 signaling, we analyzed immune populations from vehicle- and cytokine-treated mice using flow cytometry and single-cell RNA sequencing. Cells were isolated from gastric draining lymph nodes and spleens seven days after cell transfer, corresponding to six days following a single administration of vehicle or coordinated IL-2/TGF-β3 signaling. At this time point, the frequency of CD4⁺Foxp3⁺ regulatory T cells in gastric draining lymph nodes was significantly increased in treated mice (∼25%) compared with vehicle-treated controls (∼10%, Fig. 7A, B). To further define regulatory T cell phenotypes in this inflammatory setting, CD4⁺Foxp3⁺ T cells were subjected to high-dimensional flow cytometric analysis, including markers associated with regulatory activation and proliferation (Foxp3, GITR, CD25, CTLA-4, CD103, CD73, CD38, CD134, CD137, and Ki67). Unsupervised dimensionality reduction using Uniform Manifold Approximation and Projection (UMAP) revealed distinct clustering of regulatory T cells from vehicle-treated versus cytokine-treated mice, indicating divergent protein expression profiles between groups (Fig. 7C). Notably, regulatory T cells from treated mice exhibited higher expression of CD25 compared with those from vehicle-treated controls (Fig. 7D), consistent with enhanced IL-2 responsiveness in the disease setting.

**Fig. 7.**
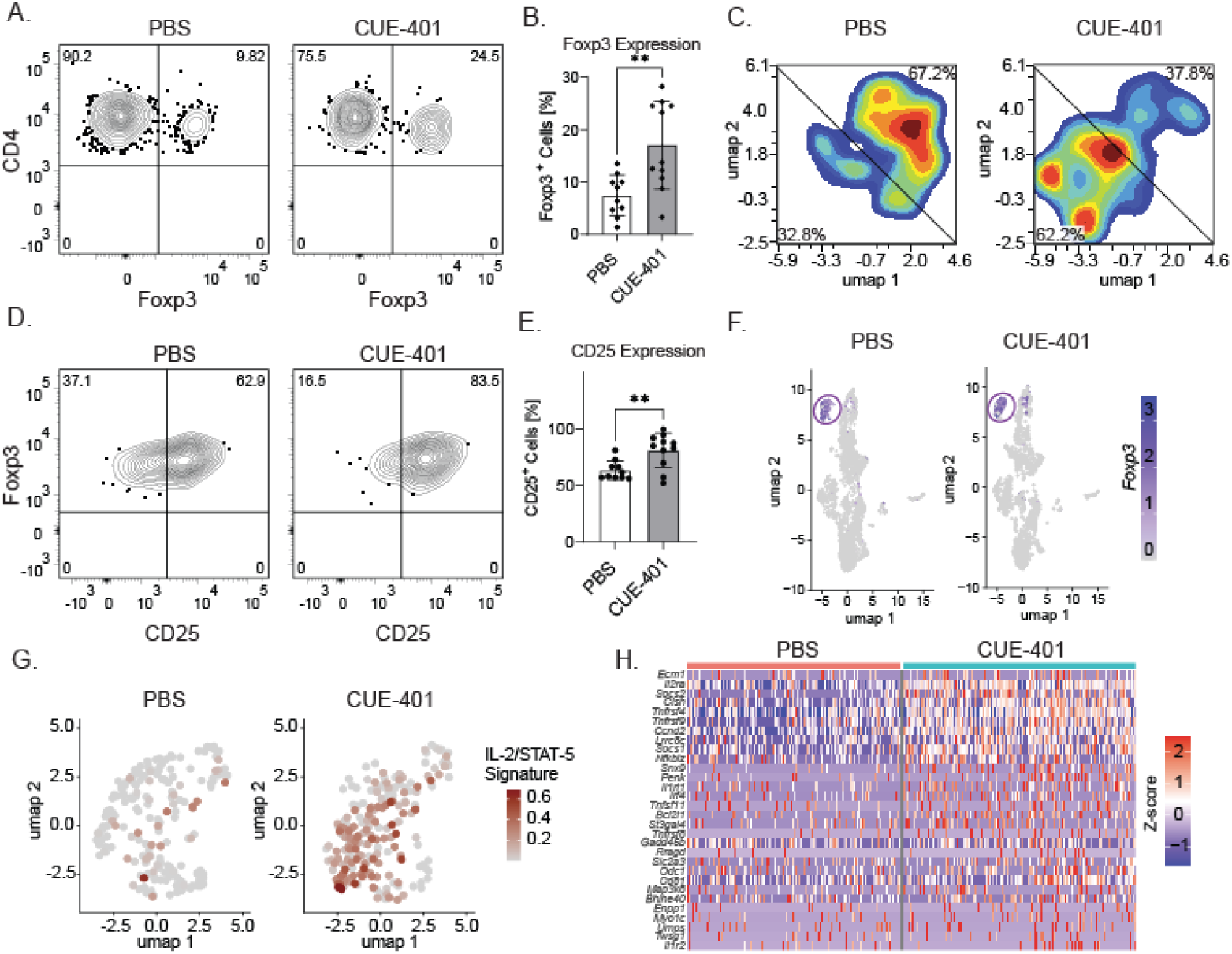
CUE-401 induces activated Tregs with a distinct transcriptional program. **(A)**. Representative flow cytometric analysis of CD4 gated Foxp3^+^ cells in PBS and CUE-401 treated BALB/c nude mice 7 days after splenocyte transfer. (**B**). Cumulative data showing the percentages of CD4^+^Foxp3^+^ Tregs in PBS and CUE-401 treated mice 6 days after a single treatment. (**C**). CD4⁺Foxp3⁺ Tregs were identified in each treatment condition, and UMAP projections generated distinct clusters of cells based on protein expression. (**D**). Representative flow cytometric analysis of CD4^+^Foxp3^+^ gated, CD25 expressing cells. (**E**). Combined data showing the percentages of CD25 expressing Tregs in PBS and CUE-401 treated mice. (**F**). Single cell RNA sequencing clustering analysis of T cells. T cell populations were subset and Foxp3 expression identified the Treg populations in PBS and CUE-401 treated mice. (**G**). Foxp3 expressing T cells were re-clustered and signatures were generated based on IL-2/STAT5 signaling transcripts that were identified through gene set enrichment analysis (**H**). Heatmap that highlights differentially expressed genes in single Foxp3^+^ cells between PBS and CUE-401 treatment. Differential expression analysis between CUE-401 and PBS Tregs used the Wilcoxon rank-sum test implemented in Seurat. P-values were adjusted for multiple comparisons using the Benjamini–Hochberg method to control false discovery rate (FDR). Representative images in (A) from 3 independent experiments with 3-5 age matched female mice per experiment. *P*-values were determined utilizing Mann-Whitney *U* test with *p* < 0.05 as significant for the flow cytometry analyses. Single cell analysis in (F-H) generated cells of 8 total mice in 2 independent experiments (4 treated and 4 untreated).

To further characterize regulatory T cell states at the transcriptional level, we performed single-cell RNA sequencing of cells isolated from gastric lymph nodes and spleens at the same time point. Foxp3⁺ regulatory T cells formed discrete clusters in each treatment condition (Fig. 7F). Downstream analysis of Foxp3⁺ cells revealed that regulatory T cells from cytokine-treated mice were significantly enriched for transcripts associated with IL-2/STAT5 signaling compared with regulatory T cells from vehicle-treated controls (Fig. 7G).

Consistent with these findings, differential gene expression analysis highlighted increased expression of genes associated with regulatory T cell activation, survival, and suppressive potential in treated mice, including *Il2ra* (CD25), *Tnfrsf4* (OX40), *Tnfrsf9* (4-1BB), *Tnfsf11* (RANKL), *Il1rl1* (ST2), and *Bcl2l1* (BCL-xL) (Fig. 7H). Together, these data indicate that coordinated IL-2 and TGF-β3 signaling in a disease context not only expands regulatory T cell populations but also promotes a distinct transcriptional and phenotypic state associated with enhanced activation and survival. These early changes in regulatory T cell state provide mechanistic insight into how regulatory dominance is established in an inflammatory environment and may contribute to subsequent long-term suppression of autoimmune pathology.

## Discussion

Regulatory T cell–mediated immune tolerance emerges from the integration of multiple cytokine-derived signals that collectively shape lineage stability, activation state, and functional dominance over effector responses. Although interleukin-2 (IL-2) and transforming growth factor-β (TGF-β) have long been recognized as essential contributors to regulatory T cell biology, how these pathways can be coordinated in vivo to selectively promote regulatory outcomes while constraining conventional T cell activation has remained incompletely defined. In this study, we demonstrate that constrained and coordinated delivery of attenuated IL-2 and TGF-β signals is sufficient to induce, expand, and stabilize regulatory T cells across multiple cellular contexts, leading to durable suppression of autoreactive immune responses.

Prior studies established that inducible regulatory T cells generated ex vivo with IL-2 and TGF-β can suppress autoimmune pathology following adoptive transfer (*33–36, 39, 40*). However, these approaches require cell manufacturing and do not address whether coordinated cytokine delivery can be achieved pharmacologically in vivo. Our findings show that coordinated delivery of IL-2 and TGF-β3 signaling can recapitulate key features of regulatory T cell induction and activation directly in vivo while avoiding broad activation of conventional CD4⁺ T cells. Notably, regulatory T cells exposed to coordinated signaling acquired an activated phenotype characterized by increased expression of suppressive and survival-associated receptors, whereas such activation was not observed in Foxp3⁻ conventional T cells. This selective response was not reproduced by an IL-2–only variant, supporting the premise that integration of TGF-β3 signaling qualitatively reshapes IL-2 responsiveness in a Treg-biased manner. Further dissection of receptor-proximal signaling dynamics and cell-type–specific cytokine engagement will be required to refine this model (*41*).

At the molecular level, regulatory T cells expanded under coordinated IL-2/TGF-β3 signaling exhibited durable Foxp3 expression, demethylation of the Foxp3 Treg-specific demethylated region (TSDR), and stable maintenance of Foxp3, all features associated with lineage commitment. These properties are particularly notable given reports that inducible regulatory T cells can exhibit instability under inflammatory conditions (*42–45*). Although overall regulatory T cell frequencies returned toward baseline approximately two weeks after treatment, expanded cells retained a stable Foxp3⁺ phenotype in both homeostatic and inflammatory settings, consistent with physiological regulation of the compartment rather than loss of lineage identity. Future lineage-tracing studies will be important to definitively distinguish contributions of de novo induction versus selective expansion in vivo.

Mechanistically, coordinated IL-2 and TGF-β signaling promoted activation of CD25/IL-2/STAT5–associated pathways in regulatory T cells while limiting induction of effector-associated transcriptional programs in conventional CD4⁺ T cells. Comparative studies using a TGF-β–deficient IL-2 variant demonstrated that TGF-β3 is required both for selective regulatory T cell activation and for constraining off-target IL-2–driven responses in conventional T cells. These findings indicate that the integration of IL-2 and TGF-β signaling does not simply amplify IL-2 activity but instead establishes a qualitatively distinct signaling state biased toward regulatory outcomes. Whether this effect reflects altered cytokine receptor geometry, localized cytokine concentration, temporal coordination of signaling pathways, or cooperative receptor engagement remains an important question for future mechanistic investigation.

The immunological relevance of this coordinated signaling logic was particularly evident in a stringent model of regulatory T cell deficiency–driven autoimmunity. Early, transient exposure to coordinated IL-2 and TGF-β signaling suppressed accumulation of autoreactive T cells, reduced tissue pathology, and promoted establishment of regulatory dominance that persisted long after cytokine exposure had ceased. Single-cell transcriptional analysis further revealed that regulatory T cells expanded under these conditions acquired a distinct activation program characterized by enhanced IL-2/STAT5 signaling, expression of costimulatory receptors, and pro-survival genes in disease-associated lymphoid tissues, suggesting that early regulatory reprogramming contributes to long-term immune control.

While this study demonstrates that coordinated IL-2 and TGF-β3 signaling can promote regulatory T cell dominance without enforced receptor co-engagement or antigen targeting, recent work by Sun et al. describes an alternative strategy in which IL-2 and helminth derived TGF-β mimic are conditionally coupled through a cis-acting surrogate agonist to drive peripheral regulatory T cell differentiation in vivo (*46*). Together, these studies define complementary mechanistic solutions by which IL-2 and TGF-β signaling can be integrated in vivo, reinforcing the concept that qualitative features of signal integration, rather than cytokine availability alone, govern regulatory T cell fate and function.

Collectively, our findings establish coordinated and attenuated IL-2 and TGF-β3 signaling as a flexible immunological framework for regulating regulatory T cell responses in vivo. By demonstrating that signal attenuation and integration can selectively promote stable, lineage-committed regulatory T cells while limiting conventional T cell activation, this work provides insight into fundamental principles governing immune tolerance. Evaluation in additional disease models and human systems will further clarify how distinct modes of cytokine coordination can be leveraged to engage regulatory pathways across diverse immune contexts.

## Materials and Methods

### Study design

The objective of this research was to define the activities of a new biologic that co-engages IL-2 and TGF-β pathways on regulatory T cells. This was accomplished by experimentally testing the function of the biologic, named CUE-401, in vitro and in vivo. In vitro, testing was performed to address the induction of iTregs from naïve CD4^+^ T cells, maintenance of Foxp3 expression and expansion of existing Tregs, and induction of Foxp3 expression with reduced proinflammatory cytokine production in a subset of effector/memory CD4⁺ T cells. CUE-401 was also administered to mice. Expression of Foxp3 and other activation related proteins were measured by flow cytometry and compared by uniform manifold approximation and projection analysis. Stability of in vivo Tregs was assessed by measuring the methylation state of CpG sites in the Treg specific demethylated region, and Foxp3^+^ cells were restimulated to verify maintenance of Foxp3 expression. CUE-401’s ability to affect disease severity was measured through tracking autoreactive T cells, pathological scores of disease in the stomachs of an established mouse model of autoimmunity. Single cell RNA sequencing was done to address potential mechanisms of action for disease.

### CUE-401 design and production

CUE-401 was produced by transient transfection of ExpiCHO-S cells using the Gibco ExpiCHO Expression System Max Titer Protocol. Each transfection is performed with two plasmids at a 1:1 molar ratio encoding (1) IL2 mutein linked to human IgG1Fc (knob) linked to monomeric TGFß3 and (2) human IgG1Fc (hole) linked to truncated TGFßR2, respectively. Upon harvest, conditioned media was loaded onto a mAb Select SuRe protein A column using an AKTA Pure 25 system for affinity-based purification. After a 10-column volume (cv) wash with 20mM HEPES pH 7.5 + 1M NaCl, CUE-401 was eluted using 50mM glycine pH 2.8 directly into 10% 1M Tris pH 9.0 neutralization solution. Eluant was pooled, concentrated and loaded onto a HiLoad 26/600 Superdex 200 for final step purification by size exclusion chromatography in 1x PBS pH 7.4. Main protein elution peak was collected, pooled, concentrated, filtered using a 2um syringe filter. Protein concentration was measured by absorbance at 280nm (A_280_) using a NanoDrop system. Protein integrity, size, solution state, and purity were confirmed by liquid chromatography-mass spectrometry, SDS-PAGE, and analytical size exclusion chromatography. Final purified protein was aliquoted and stored at −80°C until use.

### Mice

Female BALB/c Thy1.1^+^ mice were bred in house and BALB/c *nu/nu* (4–8 wk old) were purchased from the Charles River Laboratory and housed under specific pathogen-free conditions. TCR transgenic mice were crossed with BALB/c Thy1.1/1.1 expressing mice as previously described (*37*). TCR Transgenic Thy1.1/1.1 expressing mice that develop autoimmune gastritis were crossed with BALB/c Thy1.1/1.1 Foxp3.eGFP reporter mice (C.Cg-*Foxp3^tm2Tch^*/J) to generate TCR transgenic Thy1.1/1.1 Foxp3.eGFP reporter mice (*42*). All animals were maintained in the Department of Comparative Medicine, Saint Louis University School of Medicine in accordance with institutional guidelines (Association for Assessment and Accreditation of Laboratory Animal Care, International accreditation number 000656).

### CUE-401 Dosing

CUE-401 was formulated in sterile PBS, pH 7.4. Mice received intraperitoneal (i.p.) injections of CUE-401 at 2 mg/kg (100 μg/mL; 10 mL/kg; typical injection volume ∼0.2–0.3 mL per 20–30 g mouse) or PBS (vehicle) on the indicated schedule. For the adoptive transfer model, dosing occurred on days +1 and +14 post-transfer. For pharmacodynamic studies of Treg expansion, a single dose was administered, and tissues were analyzed on day 6.

### CTLL-2 proliferation assays

CTLL-2 cells were cultured in RPMI 1640 medium (Thermo Fisher Scientific, Waltham, MA, USA) supplemented with 10% heat-inactivated fetal bovine serum (Thermo Fisher Scientific, Waltham, MA, USA) and were starved of IL-2 for 22–24 h by washing cells free of cytokine and incubating them at 1×10⁵ cells/mL in IL-2–free medium at 37°C with 5% CO₂. Cell viability and morphology were assessed before and after starvation via acridine orange/propidium iodide staining using a Cellaca MX automated cell counter (Nexcelom Bioscience, Lawrence, MA, USA). On the day of assay, 5×10³ viable CTLL-2 cells were seeded in 96-well tissue-culture-treated plates (Corning/Costar, Corning, NY, USA) in 100 μL medium, with outer wells filled with DPBS (Thermo Fisher Scientific, Waltham, MA, USA) to minimize edge effects. Test molecules and recombinant rhIL-2 were prepared at 2× concentration and added in 100 μL to achieve a final volume of 200 μL per well, with all conditions plated in triplicate. After 72 h incubation at 37°C, cell viability was quantified using CellTiter-Glo^®^ reagent (Promega, Madison, WI, USA): plates were equilibrated to room temperature, 100 μL of suspension was transferred to white flat-bottom 96-well assay plates (Corning/Costar, Corning, NY, USA), an equal volume of CellTiter-Glo was added via reverse pipetting, plates were agitated and incubated for 10 min, and luminescence was measured using a Synergy Neo2 multimode plate reader (BioTek Instruments, Winooski, VT, USA).

### pSMAD Detection

Murine splenocytes and human PBMCs were used to evaluate pSMAD2 signaling in response to CUE-401 stimulation and TGF-β pathway inhibition. Fresh spleens from BALB/c mice were processed through a 0.7-µm strainer to obtain single-cell suspensions, followed by ACK lysis, washing in PBS, and counting on a Cellaca MX automated cell counter. Cells were aliquoted at 5×10⁷ per condition and rested for 2 h at 37°C, with subsets pre-treated for 90 min with 10 µM TGF-β inhibitor. PBMCs were thawed and washed similarly. Cells were treated for 30 min at 37°C with IL-2, TGF-β3, CUE-401, or combinations thereof, and then washed and lysed with RIPA buffer (Thermo Fisher Scientific, Waltham, MA, USA) supplemented with Halt Protease/Phosphatase Inhibitor Cocktail (Thermo Fisher Scientific, Waltham, MA, USA). Lysates were homogenized by pipetting, clarified by centrifugation at 10,000 rpm for 10 min at 4°C, and protein content was quantified using a BCA assay. Samples were mixed with LDS loading buffer (Thermo Fisher Scientific, Waltham, MA, USA), heated, and resolved on Bolt 4–12%

Bis-Tris Plus polyacrylamide gels (Thermo Fisher Scientific, Waltham, MA, USA) alongside SeeBlue Plus Prestained Standards (Thermo Fisher Scientific, Waltham, MA, USA). Following electrophoresis, proteins were transferred to PVDF membranes using the iBlot 2 Dry Blotting System (Thermo Fisher Scientific, Waltham, MA, USA), then blocked for 2 h at room temperature in SuperBlock buffer (Thermo Fisher Scientific, Waltham, MA, USA). Membranes were incubated overnight at 4°C with primary monoclonal antibodies against phospho-SMAD2 (Cell Signaling Technology, Danvers, MA, USA) or total SMAD2/3 (Cell Signaling Technology, Danvers, MA, USA). After washing, membranes were incubated with HRP-conjugated goat anti-rabbit IgG (Cell Signaling Technology, Danvers, MA, USA) for 1 h at room temperature, washed again, and developed using SuperSignal West Femto or West Pico chemiluminescent substrates (Thermo Fisher Scientific, Waltham, MA, USA). Western blots were imaged using a Syngene G:BOX darkroom imaging system (Syngene, Frederick, MD, USA). Band intensities were analyzed to quantify pSMAD2 induction and inhibition across treatment conditions.

### Isolation of Immune Cells

Isolation of immune cells from stomach tissue has been described previously (*38, 43*). Briefly, stomachs were harvested from mice, gastric lymph nodes were removed, and the stomachs were dissected along the lesser curvature. Stomachs were rinsed with cold PBS then injected with 10 mL of cold MACS buffer (1 x PBS,0.5% BSA (Fisher Scientific Waltham, MA), 2 mM EDTA (Promega, Fitchburg, Wisconsin)) using a 27-gauge needle, to inflate the tissue and free infiltrating immune cells. After injection, the stomach was diced and collected with the 10mL of MACS buffer and transferred to a 50 mL conical tube. The dispersed stomach tissue was mixed thoroughly then passed through a 40-micron filter, washed with 40 mL of MACS buffer and used for analysis. Immune cells from the spleen and gastric/pancreatic lymph node were processed by dispersing on a 40-micron filter and washed with 15 mL of MACS buffer. Spleen cells were spun down at 2000 RPM for 5 minutes after initial processing. Spleen cells were resuspended in 5 mL of 1x ACK lysis buffer (Thermo Fisher Scientific, Waltham, MA) for 10 seconds then washed with 45 mL of MACS buffer and spun again at 2000 RPM for 5 minutes. Cells were resuspended and filtered again through a 40-micron filter before being counted and utilized for experiments.

### T cell purification, iTreg differentiation, and nTreg expansion

Inducible Tregs (iTreg) were generated as previously described (*9*). Briefly, CD4^+^CD8^-^ thymocytes or CD4^+^Foxp3.eGFP^+^ splenocytes were sorted from 6-8-wk-old BALB/c mice and stimulated with plate-bound anti-CD3 (1 mg/ml) and anti-CD28 (2 mg/ml; BD Biosciences) in 24-well plates (2.5×10^5^ cells/well) in 2 ml RPMI media (modified RPMI 1640 (Corning, Tewksbury, Massachusetts) supplemented with 10% fetal bovine serum (FBS; Biotechne, Minneapolis, Minnesota), 2 mM glutamine (Sigma-Aldrich), 100 U/ml penicillin,100 mg/ml streptomycin (Sigma-Aldrich), 10 mM HEPES (Sigma-Aldrich), β-mercaptoethanol (55mM), 1 mM essential amino acids (Gibco, Billings, Montana), and 1 mM sodium pyruvate (Sigma-Aldrich) supplemented with recombinant human IL-2(rhIL-2) (National Institutes of Health, Bethesda, Maryland) (100 U/ml). Recombinant human TGF-β3 (rhTGF-β3) (Biotechne) (5 ng/ml) was added to induce iTregs. Cells were removed from plate-bound antibodies after 48 hours and added to new wells with fresh media supplemented with 100 U/ml recombinant human IL-2. Cells were used after 7 days in culture. Endogenous Treg expansion was assessed by isolating FACS-purified splenic CD4⁺Foxp3.eGFP⁺ Tregs. Tregs were stimulated with plate-bound anti-CD3 & anti-CD28 and expanded with IL-2 (100 U/mL) or CUE-401 as indicated for 5–7 days. Foxp3 maintenance and cell counts were assessed by flow cytometry.

### Effector/Memory CD4⁺ T-Cell Restimulation and Cytokine Quantification

Splenic CD4⁺CD44^hi^ Foxp3.eGFP⁻ T cells were FACS-sorted and stimulated with plate-bound anti-CD3 & anti-CD28 in the presence of rhIL-2 (100 U/mL), rhIL-2 + rhTGF-β3 (5 ng/mL), or CUE-401. For cytokine secretion, cells were cultured at 1×10⁶ cells/mL and restimulated with 25ng/mL phorbol myristate acetate (PMA) (Sigma-Aldrich, St. Louis, Missouri), and 500 ng/mL ionomycin (Sigma-Aldrich), for 72 hours at 37°C for 72 h. Supernatants were collected and cytokine secretion was analyzed using a Th1, Th2, Th17 and IL-13 flex set (BD Biosciences) cytometric bead array kit and analyzed by flow cytometry following the kit protocol and analyzed by flow cytometry.

### Treg Stability: TSDR (CNS2) Methylation Analysis

CD4⁺Foxp3⁺ and CD4⁺Foxp3⁻ cells were sorted from spleens of PBS- or CUE-401-treated mice 6 days post-dose. Genomic DNA was isolated (DNeasy Blood & Tissue Kit, Qiagen), bisulfite-converted (EZ DNA Methylation-Gold Kit, Zymo Research), and the Foxp3 TSDR (CNS2) region was PCR-amplified using validated primers spanning CpG sites within the enhancer. Amplicons were analyzed by pyrosequencing [or Sanger sequencing of cloned products] to quantify methylation at individual CpGs. Percent demethylation was calculated per CpG and averaged across the TSDR. Males and female mice were used for TSDR analyses. Methylation state was normalized in female mice to account for the X-linked Foxp3 locus inactivation in females.

### Bulk RNA-seq (Tregs and Tconv)

CD4⁺Foxp3.eGFP^+^ regulatory T cells (Tregs) and CD4⁺Foxp3.eGFP^-^ conventional T cells (Tconv) were FACS-sorted from spleens of PBS-, CUE-401–, or CUE-401ΔTGF-β3–treated BALB/c mice 6 days after treatment (≥95% purity). Sorted cells were pelleted and immediately lysed in lysis buffer supplied with the PureLink™ RNA Micro Kit (Invitrogen, Thermo Fisher Scientific), and total RNA was extracted according to the manufacturer’s protocol. RNA was eluted in RNase-free water and quantified using a Qubit RNA HS Assay (Thermo Fisher). RNA integrity was assessed using the Agilent Bioanalyzer RNA Pico Kit, and only samples with RIN ≥7.0 were used for library preparation.

Libraries were generated using the Takara-Clontech SMARTer construction method for Illumina with poly-A mRNA enrichment, following manufacturer instructions. Indexed libraries were pooled and sequenced on an Illumina NovaSeq x Pluse (300 cycles) platform (paired-end 2×100 bp), targeting ≥30 million read pairs per sample.

Samples were processed and sequenced at the Genome Technology Access Center @ McDonnell Genome Institute in St. Louis, Missouri. RNA-seq reads were then aligned to the Ensembl GRCm39.113 primary assembly with STAR version 2.7.11b. Gene counts were derived from the number of uniquely aligned fragments by Subread:featureCount version 2.0.8. Differential gene expression analysis was performed with EdgeR using quasi-likelihood F-test and was adjusted using the Benjamini-Hochberg method. Genes with a false discovery rate (FDR) < 0.05 were considered significantly differentially expressed and adjusted *P* value was p < 0.05. Level counts were obtained with featureCount. Visualization of the heatmaps was generated using the R package pheatmap. Level counts were obtained with featureCounts. Differential gene expression analyses were performed using DESeq2. Significance thresholds were set at an adjusted

### Cell purification and injection for adoptive transfer model of autoimmune gastritis

Naive TxA23 T cells were isolated from a single-cell suspension of thymii from TxA23.Thy-1.1 Foxp3.eGFP mice. Thymocytes were lysed with ACK lysis buffer (BioSource) and washed with MACS buffer prior to staining with CD4 BV421 (BD Biosciences) and CD8 PE (BD Biosciences). Cells were placed in FACS sort buffer (2% FBS (Biotechne, Minneapolis, Minnesota), 100 U/ml penicillin, 100 mg/ml streptomycin, in complete RPMI without phenol red (Sigma-Aldrich), and then purified using the BD FACSAria Fusion cell sorter (BD Biosciences) to obtain CD4^+^CD8^-^ GFP^-^ Cells. BALB/c Splenocytes were from spleens of healthy Thy-1.2 expressing BALB/c mice, RBC were removed with ACK lysis buffer and then purified as follows. To deplete CD25^+^ cells, cells were incubated for 10 min with anti-CD25-PE (PC61), washed in MACS buffer, and incubated for an additional 10 min with anti-PE MACS microbeads (Miltenyi Biotec, Bergisch Gladbach) then subjected to autoMACS purification (Miltenyi Biotec). The negative fraction was used as CD25-depleted spleen cells. A typical depletion resulted in <1% of the CD4^+^ cells expressing CD25. Prior to injection, all cells were washed twice in PBS. Naive TxA23.Thy-1.1 T cells that were isolated from thymus, were mixed into the BALB/c Thy1.2 splenocyte population such that an i.p. injection of 0.3 ml/mouse resulted in the transfer of 100,000 TxA23.Thy-1.1 T cells and 20×10^6^ spleen cells. Mice were monitored weekly and euthanized at ∼ day 62 for tissue analyses.

### Histopathology

BALB/c *nu/nu* mice were euthanized 62 days after receiving an adoptive transfer of splenocytes and naïve CD4^+^ T cells from autoreactive mice. The stomach opened, washed with cold PBS, and fixed in 4% neutral buffered formalin. Fixed tissues were embedded in paraffin, stained with hematoxylin and eosin (H&E) by Saint Louis University Research Histology Facility (SLU ASBRH) and disease pathology was scored similarly to previously published assessments (*38*). Briefly multiple pathological features, including inflammation, glandular atrophy, and epithelial hyperplasia were graded on a scale of 1–4, depending on the extent of mononuclear cell infiltration and parietal and chief cell destruction. All stomach sections were scored blindly by at least two individuals. General descriptions for scores are as follows: 0, healthy tissue with no abnormalities 0.5, scattered lymphocytes throughout submucosa and muscularis with one or two small dense blankets of lymphocytes; 1, two to four small dense clusters of lymphocytes in the submucosa/mucosa; 1.5, two or three areas with intermediate infiltration spanning one-third of mucosa; 2, big nodules of lymphocytic accumulation spanning half to all of mucosa; 2.5, big nodules of lymphocytic accumulation spanning half to all of mucosa beginning to show evidence of parietal and chief cell destruction (<25%); 3.0, heavy lymphocytic infiltration throughout mucosa, parietal and chief cell loss (25–75%), and replacement by foamy cells; and 4.0, total parietal and chief cell loss, no mucosal architecture, and many foamy cells.

### Flow Cytometry

Immune cells were isolated from gastric mucosa, gastric/pancreatic or inguinal lymph nodes, and spleens following the protocol previously described and cell surface staining was performed according to standard procedures (*38, 43*). Immune cell subsets and proteins expressed on Tregs were identified using the following antibodies: CD4 BV421, CD8 BV510, CD90.2 BV786, CD4 BV711, CD38 BV711, CD137 BV650 CD152 (CTLA4), CD103 PE-Cy7, CD8 PE, CD19 PE (BD Biosciences, Franklin Lakes, New Jersey), CD73 PE/Dazzle594, CD90.1, CD134 PerCP/Cyanine 5.5, Ki67 BV605 (BioLegend, San Diego, California), CD25 APC-eFluor780, Foxp3 APC, CD62L eFlour450, CD44 APC-eFlour780 (Thermo Fisher Scientific). Cells were washed with 200 uL of PBS and stained in only PBS at 4°C. Cells were fixed and permeabilized using eBioscience Foxp3/Transcription Factor Staining Buffer Set (Thermo Fisher Scientific). Cells were incubated overnight with the intracellular antibodies and then washed and analyzed by flow cytometry. Flow cytometry was performed on a BD LSRII, a BD Fortessa (BD Biosciences), or a Cytek Aurora (Cytek Biosciences, Fremont California), and analyzed using FlowJo (FlowJo, Ashland, OR).

### Single-Cell RNA-seq

Cells from gastric lymph nodes and spleens (day 7 post transfer; day 6 post-treatment) were prepared as above, RBC depleted as needed and filtered through 40μm strainers. Viability (>85%) was confirmed by trypan blue. Cells were loaded on a 10x Genomics Chromium Controller (10X Genomics, Pleasanton, CA) targeting up to 10,000 cells per library using the Next GEM Single Cell 5′ Kit v2 (10x #1000263). cDNA amplification and library prep followed manufacturer protocols. PCR reactions were conducted in the T100 Thermal Cycler (BioRad, Hercules, CA). Libraries were QC’d on an Agilent 2100 Bioanalyzer (Agilent, Santa Clara, CA) and sequenced on an Illumina NovaSeq 6000 (Illumina, San Diego, CA).

Raw data were processed through CellRanger 6.0.2 pipeline (10x Genomics). Conditional repeats were combined using cellranger aggr. Analysis was conducted using Seurat. (*44*) Each dataset was filtered to exclude cells with low gene expression (<500/cell), high gene counts (outliers), and high percent mitochondrial transcript load (>10%/cell). Datasets were merged, normalized, scaled, and dimensionally reduced. Layers were joined and clusters were generated using a resolution of 2. Cluster identities were established by attaining distinguishing genes per cluster and relating them to known profiles for immune cells present in spleen and gastric lymph nodes. T cells clusters were subset and pathway analysis was conducted using the R package enrichR and the Msig database for transcripts. Gene ontology, pathway enrichment, and signature analyses (e.g., IL2/STAT5 signaling) were conducted using fgsea and enrichR with MSigDB gene sets. Visualization of feature plots was performed using the ggplot2 package in combination with the scCustomize package. Heatmaps were generated using Seurat’s DoHeatMap function. Statistical analyses were performed in Prism 10.0.0 (GraphPad Software). *P* values less than 0.05 were considered significant.

### Statistical Analysis

In vitro data are expressed as means of individual determination ± SD. In vivo data are expressed as median of individual determination ± 95% confidence interval. Statistical analysis was performed using GraphPad Prism 10 (GraphPad Software, San Diego, CA). Dose titrations of CUE-401 on nTregs utilized Kruskal-Wallis test. Cytokine analysis utilized Wilcoxon matched-pairs signed rank test. In vitro treatment comparisons between rhIL-2, rhIL-2 + TGF-β, and CUE-401 utilized Ordinary one-way ANOVA with Tukey’s multiple comparisons test. Histopathology score significance was performed using the Mann–Whitney *U* test. Differential expression analysis between CUE-401 and PBS treated Tregs in scRNA-seq data was performed using the Wilcoxon rank-sum test implemented in Seurat. Differentially expressed genes were identified using the FindMarkers function in the Seurat package with default statistical parameters, a log fold-change threshold of 0.6, and a minimum expression fraction of 0.1. Genes with an adjusted p-value < 0.05 were considered differentially expressed. For bulk RNA sequencing, P-values were adjusted for multiple comparisons using the Benjamini–Hochberg method to control false discovery rate.

## Acknowledgments

We thank Saint Louis University Flow Cytometry Core (Joy Eslick, and Jacqueline Spencer) and the Advanced Research Histology Facility (Caroline Murphy, Vasiliki Grammatopoulou) for assistance with data collection. We thank Saint Louis University Genomics and Bioinformatics Core Facility (Dr. Michelle Brennan) for assistance with processing and analysis of single cell and bulk RNA sequencing. We thank Saint Louis University Department of Comparative Medicine (Lisa Tempel, Tawanika Williams, Kelsey Ernst) for assistance in animal care and handling

## Author contributions

Conceptualization: NMJ, RS, NMG, SNQ, AS, CAP, SGH, EF, RJD Methodology: NMJ, NMG, SNQ, RJD, SL, EC, KY, EN

Investigation: NMJ

Visualization: NMJ, KF, JAC, RJD

Formal Analysis: NMJ, NY, CAP, SGH, KF, EF Software: SGH, KF

Funding acquisition: AS, NMG, SNQ, RJD Writing – original draft: NMJ, JAC, RJD

Writing – review & editing: NMJ, JAC, NMG, SNQ, RJD All authors approved the final manuscript.

## Data and materials availability

All data necessary to understand and evaluate the conclusions of the paper are provided in the manuscript and supplementary materials. The CUE-401 biologic can be available upon request under a material transfer agreement with CUE Biopharma. Sequencing data will be made publicly available upon acceptance of this publication.

## References

1. S. Sakaguchi, N. Sakaguchi, M. Asano, M. Itoh, M. Toda, Immunologic self-tolerance maintained by activated T cells expressing IL-2 receptor alpha-chains (CD25). Breakdown of a single mechanism of self-tolerance causes various autoimmune diseases. J Immunol 155, 1151–1164 (1995).

2. L. Wang et al., Regulatory T cells in homeostasis and disease: molecular mechanisms and therapeutic potential. Signal Transduct Target Ther 10, 345 (2025).

3. C. L. Bennett et al., The immune dysregulation, polyendocrinopathy, enteropathy, X-linked syndrome (IPEX) is caused by mutations of FOXP3. Nat Genet 27, 20–21 (2001).

4. S. Hori, T. Nomura, S. Sakaguchi, Control of regulatory T cell development by the transcription factor Foxp3. Science 299, 1057–1061 (2003).

5. R. Khattri, T. Cox, S. A. Yasayko, F. Ramsdell, An essential role for Scurfin in CD4+CD25+ T regulatory cells. Nat Immunol 4, 337–342 (2003).

6. J. D. Fontenot, M. A. Gavin, A. Y. Rudensky, Foxp3 programs the development and function of CD4+CD25+ regulatory T cells. Nat Immunol 4, 330–336 (2003).

7. E. M. Shevach, A. M. Thornton, tTregs, pTregs, and iTregs: similarities and differences. Immunol Rev 259, 88–102 (2014).

8. A. Jones, D. Hawiger, Peripherally Induced Regulatory T Cells: Recruited Protectors of the Central Nervous System against Autoimmune Neuroinflammation. Front Immunol 8, 532 (2017).

9. T. S. Davidson, R. J. DiPaolo, J. Andersson, E. M. Shevach, Cutting Edge: IL-2 is essential for TGF-beta-mediated induction of Foxp3+ T regulatory cells. J Immunol 178, 4022–4026 (2007).

10. W. Chen et al., Conversion of peripheral CD4+CD25- naive T cells to CD4+CD25+ regulatory T cells by TGF-beta induction of transcription factor Foxp3. J Exp Med 198, 1875–1886 (2003).

11. T. L. Nguyen, N. T. Makhlouf, B. A. Anthony, R. M. Teague, R. J. DiPaolo, In vitro induced regulatory T cells are unique from endogenous regulatory T cells and effective at suppressing late stages of ongoing autoimmunity. PLoS One 9, e104698 (2014).

12. S. E. Weber et al., Adaptive islet-specific regulatory CD4 T cells control autoimmune diabetes and mediate the disappearance of pathogenic Th1 cells in vivo. J Immunol 176, 4730–4739 (2006).

13. R. J. DiPaolo et al., Autoantigen-specific TGFbeta-induced Foxp3+ regulatory T cells prevent autoimmunity by inhibiting dendritic cells from activating autoreactive T cells. J Immunol 179, 4685–4693 (2007).

14. S. G. Zheng et al., CD4+ and CD8+ regulatory T cells generated ex vivo with IL-2 and TGF-beta suppress a stimulatory graft-versus-host disease with a lupus-like syndrome. J Immunol 172, 1531–1539 (2004).

15. M. E. Raeber, D. Sahin, U. Karakus, O. Boyman, A systematic review of interleukin-2-based immunotherapies in clinical trials for cancer and autoimmune diseases. EBioMedicine 90, 104539 (2023).

16. R. Zhang et al., Low-dose IL-2 therapy in autoimmune diseases: An update review. Int Rev Immunol 43, 113–137 (2024).

17. L. Khoryati, et al., An IL-2 mutein engineered to promote expansion of regulatory T cells arrests ongoing autoimmunity in mice. Sci Immunol 5, (2020).

18. S. N. Quayle et al., CUE-101, a Novel E7-pHLA-IL2-Fc Fusion Protein, Enhances Tumor Antigen-Specific T-Cell Activation for the Treatment of HPV16-Driven Malignancies. Clin Cancer Res 26, 1953–1964 (2020).

19. D. J. Stauber, E. W. Debler, P. A. Horton, K. A. Smith, I. A. Wilson, Crystal structure of the IL-2 signaling complex: paradigm for a heterotrimeric cytokine receptor. Proc Natl Acad Sci U S A 103, 2788–2793 (2006).

20. K. M. Heaton, G. Ju, E. A. Grimm, Human interleukin 2 analogues that preferentially bind the intermediate-affinity interleukin 2 receptor lead to reduced secondary cytokine secretion: implications for the use of these interleukin 2 analogues in cancer immunotherapy. Cancer Res 53, 2597–2602 (1993).

21. T. R. Mosmann et al., Species-specificity of T cell stimulating activities of IL 2 and BSF-1 (IL 4): comparison of normal and recombinant, mouse and human IL 2 and BSF-1 (IL 4). J Immunol 138, 1813–1816 (1987).

22. K. Abnaof et al., TGF-beta stimulation in human and murine cells reveals commonly affected biological processes and pathways at transcription level. BMC Syst Biol 8, 55 (2014).

23. J. E. Zuniga et al., Assembly of TbetaRI:TbetaRII:TGFbeta ternary complex in vitro with receptor extracellular domains is cooperative and isoform-dependent. J Mol Biol 354, 1052–1068 (2005).

24. J. Groppe et al., Cooperative assembly of TGF-beta superfamily signaling complexes is mediated by two disparate mechanisms and distinct modes of receptor binding. Mol Cell 29, 157–168 (2008).

25. X. Li, Y. Liang, M. LeBlanc, C. Benner, Y. Zheng, Function of a Foxp3 cis-element in protecting regulatory T cell identity. Cell 158, 734–748 (2014).

26. A. S. Wegrzyn, A. E. Kedzierska, A. Obojski, Identification and classification of distinct surface markers of T regulatory cells. Front Immunol 13, 1055805 (2022).

27. C. Gootjes, J. J. Zwaginga, B. O. Roep, T. Nikolic, Defining Human Regulatory T Cells beyond FOXP3: The Need to Combine Phenotype with Function. Cells 13, (2024).

28. S. Dikiy, A. Y. Rudensky, Principles of regulatory T cell function. Immunity 56, 240–255 (2023).

29. J. D. Fontenot, J. P. Rasmussen, M. A. Gavin, A. Y. Rudensky, A function for interleukin 2 in Foxp3-expressing regulatory T cells. Nat Immunol 6, 1142–1151 (2005).

30. A. Arvey et al., Inflammation-induced repression of chromatin bound by the transcription factor Foxp3 in regulatory T cells. Nat Immunol 15, 580–587 (2014).

31. J. A. Hill et al., Foxp3 transcription-factor-dependent and -independent regulation of the regulatory T cell transcriptional signature. Immunity 27, 786–800 (2007).

32. N. Sugimoto et al., Foxp3-dependent and -independent molecules specific for CD25+CD4+ natural regulatory T cells revealed by DNA microarray analysis. Int Immunol 18, 1197–1209 (2006).

33. M. Ahn, J. Dostal, P. Hegde, D. J. Lee, CTLA-4 expression on tregs is needed for suppression of autoimmune uveitis. Sci Rep 15, 17745 (2025).

34. M. G. Petrillo et al., GITR+ regulatory T cells in the treatment of autoimmune diseases. Autoimmun Rev 14, 117–126 (2015).

35. S. Tagkareli et al., CD103 integrin identifies a high IL-10-producing FoxP3(+) regulatory T-cell population suppressing allergic airway inflammation. Allergy 77, 1150–1164 (2022).

36. K. E. Earle et al., In vitro expanded human CD4+CD25+ regulatory T cells suppress effector T cell proliferation. Clin Immunol 115, 3–9 (2005).

37. R. S. McHugh, E. M. Shevach, D. H. Margulies, K. Natarajan, A T cell receptor transgenic model of severe, spontaneous organ-specific autoimmunity. Eur J Immunol 31, 2094–2103 (2001).

38. R. J. DiPaolo, D. D. Glass, K. E. Bijwaard, E. M. Shevach, CD4+CD25+ T cells prevent the development of organ-specific autoimmune disease by inhibiting the differentiation of autoreactive effector T cells. J Immunol 175, 7135–7142 (2005).

39. S. Gaggero, et al., IL-2 is inactivated by the acidic pH environment of tumors enabling engineering of a pH-selective mutein. Sci Immunol 7, eade5686 (2022).

40. W. J. Leonard, J. X. Lin, Strategies to therapeutically modulate cytokine action. Nat Rev Drug Discov 22, 827–854 (2023).

41. D. A. Horwitz et al., Nanoparticles loaded with IL-2 and TGF-beta promote transplantation tolerance to alloantigen. Front Immunol 15, 1429335 (2024).

42. W. Lin et al., Regulatory T cell development in the absence of functional Foxp3. Nat Immunol 8, 359–368 (2007).

43. F. Alderuccio, B. H. Toh, P. A. Gleeson, I. R. van Driel, A novel method for isolating mononuclear cells from the stomachs of mice with experimental autoimmune gastritis. Autoimmunity 21, 215–221 (1995).

44. Y. Hao et al., Dictionary learning for integrative, multimodal and scalable single-cell analysis. Nat Biotechnol 42, 293–304 (2024).

45. D. Haribhai et al., Regulatory T cells dynamically control the primary immune response to foreign antigen. J Immunol 178, 2961–2972 (2007).

46. Q. Sun et al., Facile induction of immune tolerance by an interleukin-2-TGFbeta surrogate agonist. Nature, (2026).

